## Supplemental Figures for "The evolution of transposable elements in *Brachypodium distachyon* is governed by purifying selection, while neutral and adaptive processes play a minor role"

**Supplementary Figures**

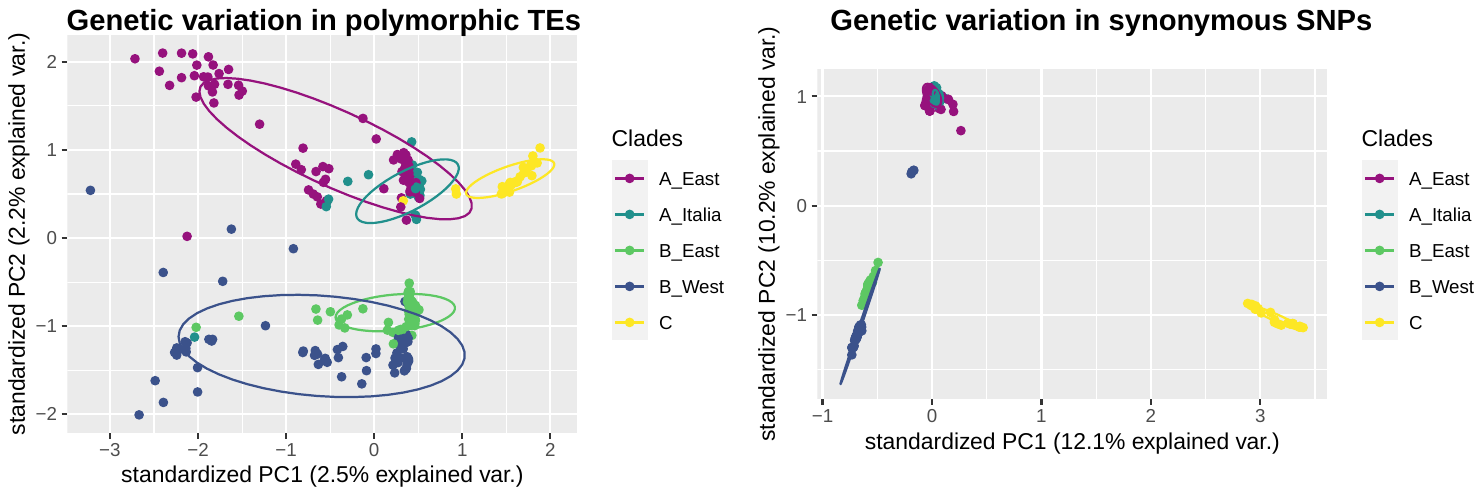

**Supplementary Figure S1.** Principal Component Analyses using TE (left panel) and SNP (right panel) polymorphisms.

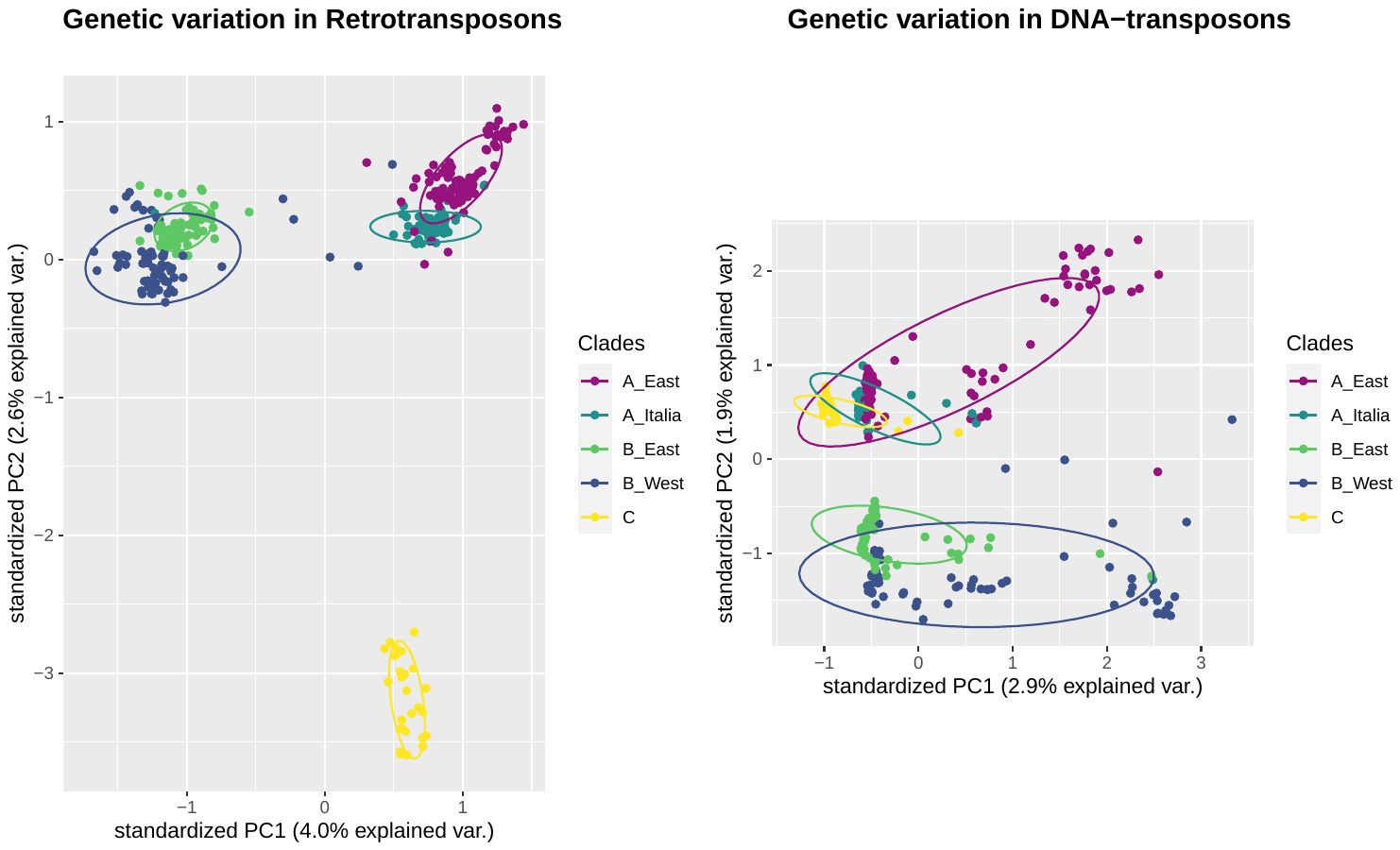

**Supplementary Figure S2.** Principal Component Analyses using retrotransposon (left panel) and DNA-transposon (right panel) polymorphisms.

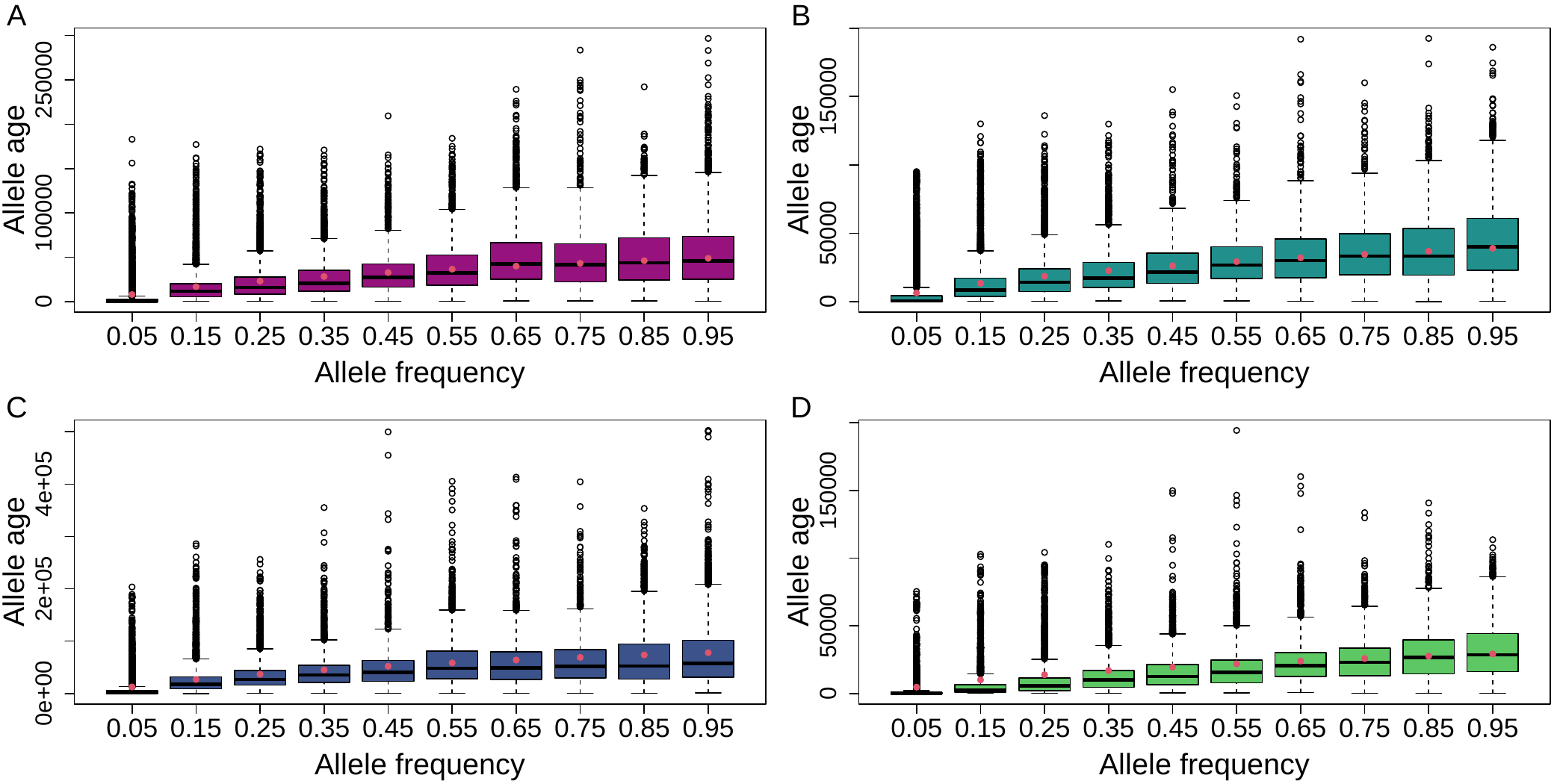

**Supplementary Figure S3.** Observed correlation between age in generations and frequency of synonymous SNPs in the four derived genetic clades. The red points show the expected age of a neutrally evolving mutation at a specific frequency based on the predictions of Kimura and Ohta (1973). Panel A: clade A_East; panel B: clade A_Italia; panel C: clade B_West and panel D: clade B_East.

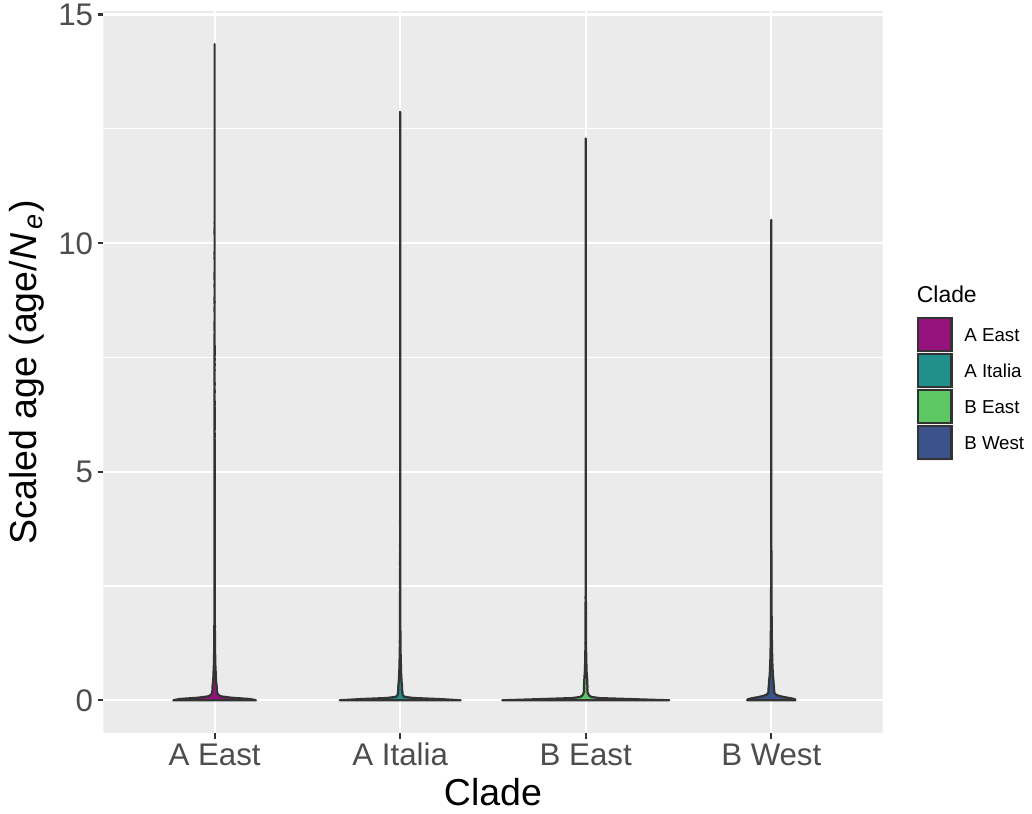

**Supplementary Figure S4.** Distribution of the observed TE age scaled by the effective population size (*N_e_*) in the four derived genetic clades of *B. distachyon*. The age estimates were scaled by the effective population size to improve readability.

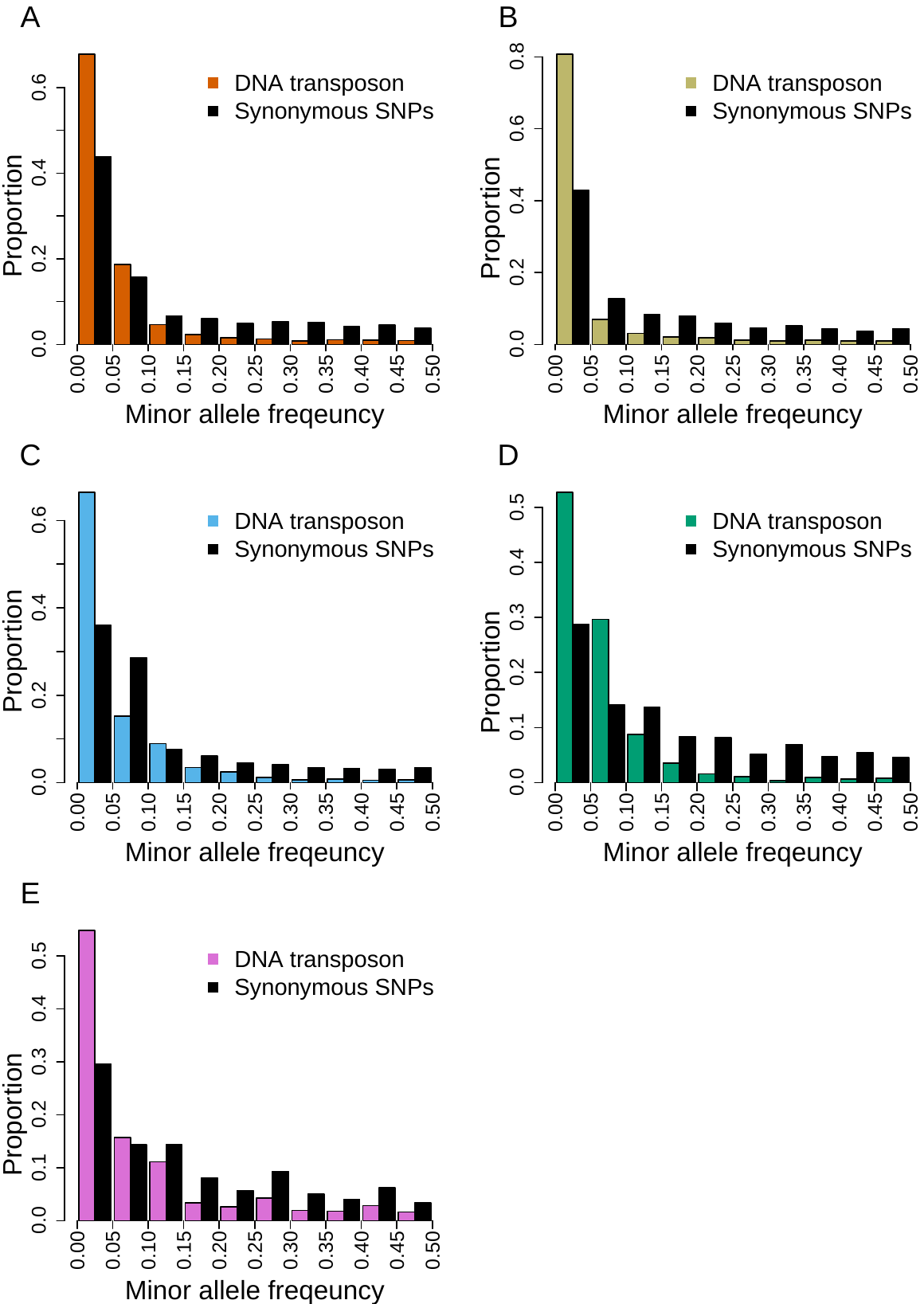

**Supplementary Figure S5.** Folded site frequency spectrum of DNA-transposons and synonymous SNPs in all genetic clades. Panel A: A_East; panel B: A_Italia; panel C: B_West; panel D: B_East; panel E: C.

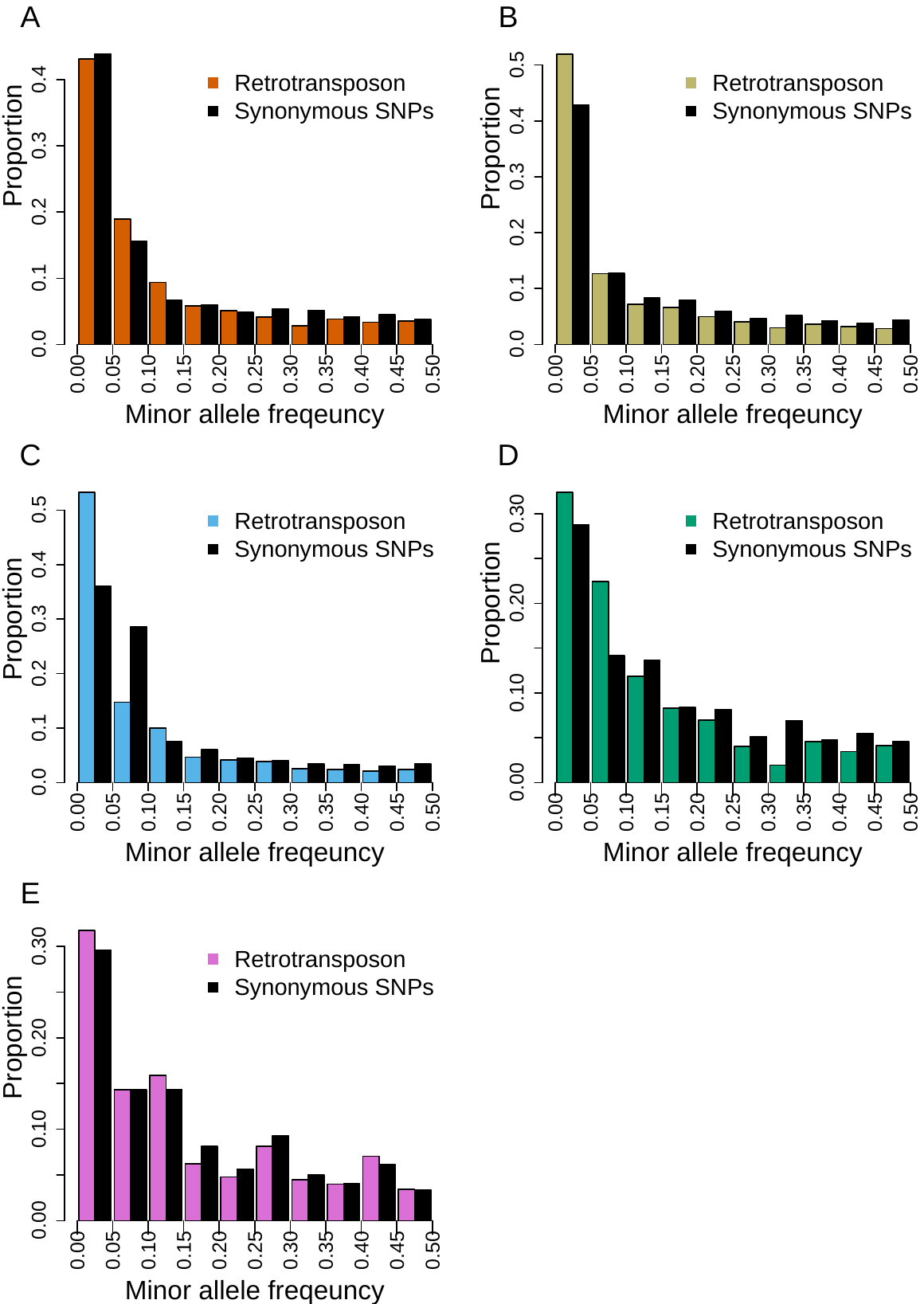

**Supplementary Figure S6.** Folded site frequency spectrum of retrotransposons and synonymous SNPs in all genetic clades. Panel A: A_East; panel B: A_Italia; panel C: B_West; panel D: B_East; panel E: C.

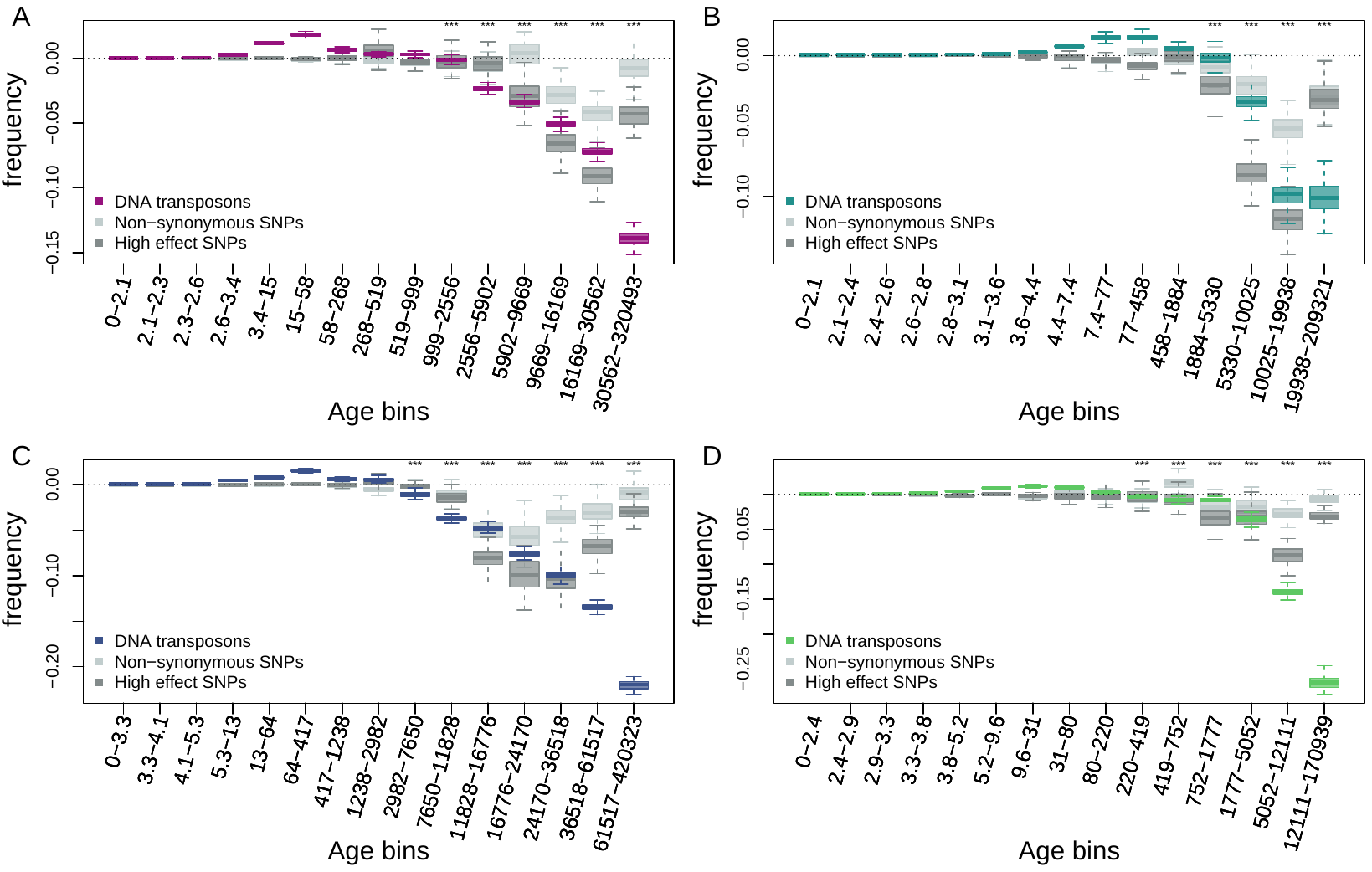

**Supplementary Figure S7.** Age-adjusted SFS of DNA-transposons (colored), non-synonymous SNPs (light gray) and high effect SNPs (dark gray) in the four derived clades. The X axes show the age range of the mutations in each bin, and the age range of each bin was chosen so that each bin represents the same number of DNA-transposons observations. A: A_East genetic clades; B: A_Italia genetic clades; C: B_West genetic clades; D: B_East genetic clades. Boxplots are based on 100 estimations of Δ frequency. Significant deviations of Δ frequency estimates from 0 in the age-adjusted SFS of DNA-transposons are shown with asterisks (one-side Wilcoxon tests, Bonferroni corrected p value < 0.01: ***).

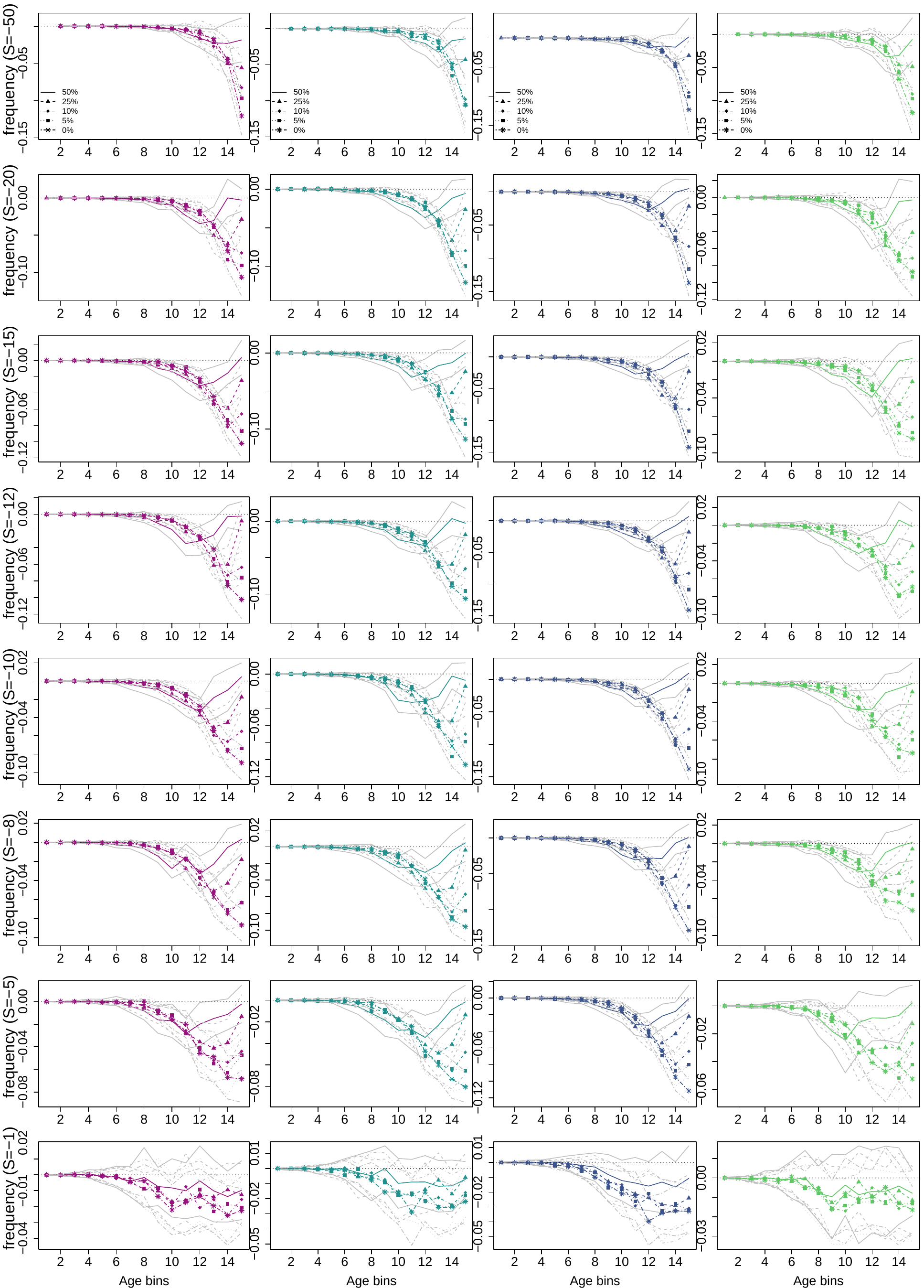

**Supplementary Figure S8.** Age-adjusted SFS of simulated mutations under negative selection in the four derived clades. The four columns show the results for the A_East, A_Italia, B_West and B_East genetic clades, respectively. Each line shows the results for the different scaled selection coefficients (S). The five colored curves in each plot show the shape of the age-adjusted SFS with varying ratios of neutrally evolving mutations, and the gray curves show variation within one standard deviation based on the 20 runs for each simulation. The X axes show the age bin from the youngest to the oldest, with each age bin including the same number of observations for each simulation.

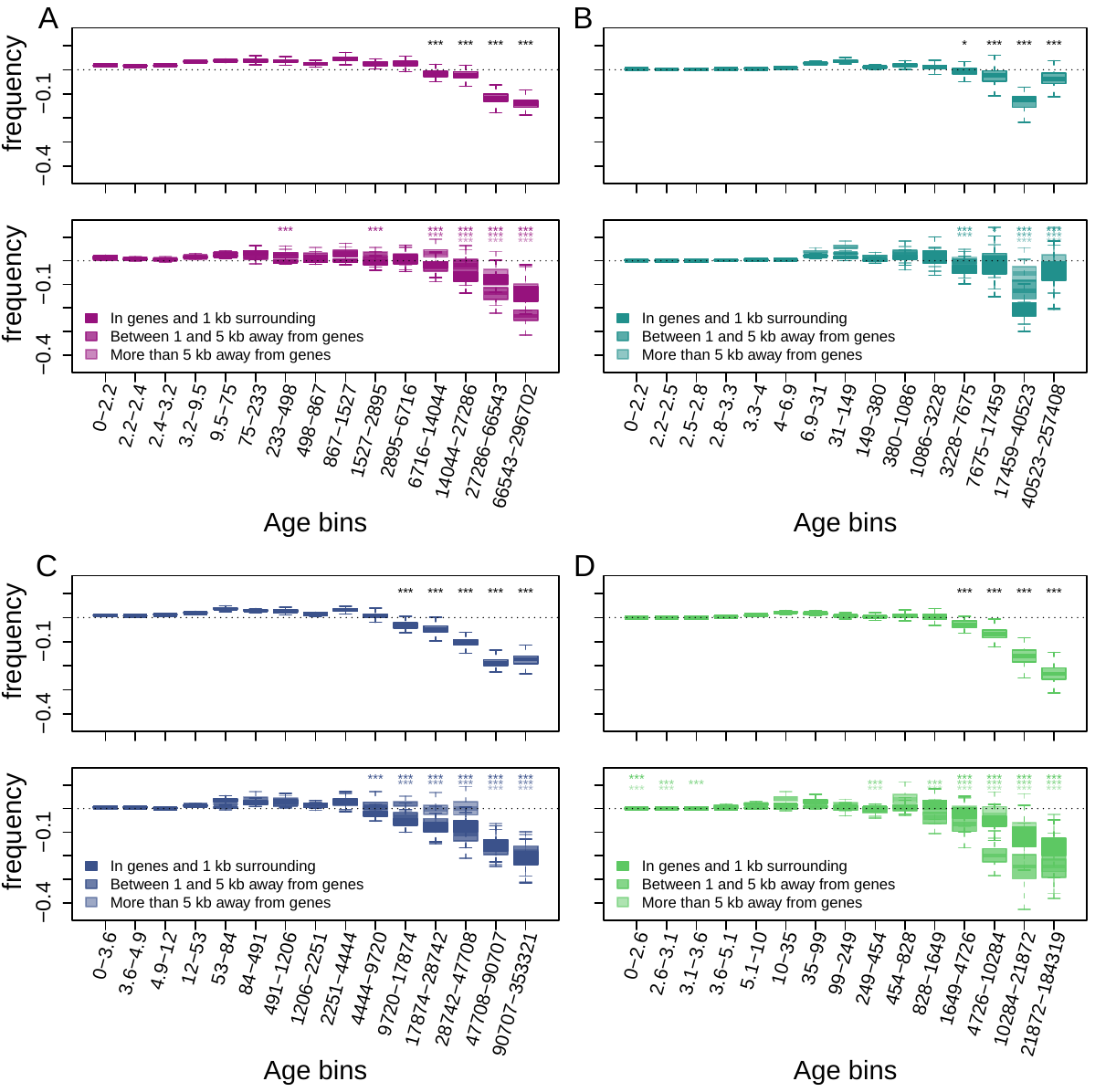

**Supplementary Figure S9.** Age-adjusted SFS of retrotransposons in accessions with at least 20x coverage. The top row shows the age-adjusted SFS of retrotransposons in the four derived clades. The bottom row shows the age-adjusted SFS of retrotransposons based on their distance to the next gene in the four derived clades. The X axes show the age range of the mutations in each bin, and the age range of each bin was chosen so that each bin represents the same number of retrotransposon observations in the top row. The different columns show the four derived clades: A: A_East; B: A_Italia; C: B_West; D: B_East. Boxplots are based on 100 estimations of Δ frequency. Significant deviations of Δ frequency estimates from 0 in the age-adjusted SFS of retrotransposons are shown with asterisks (one-side Wilcoxon tests, Bonferroni corrected *p* value < 0.05: *; < 0.01: ***).

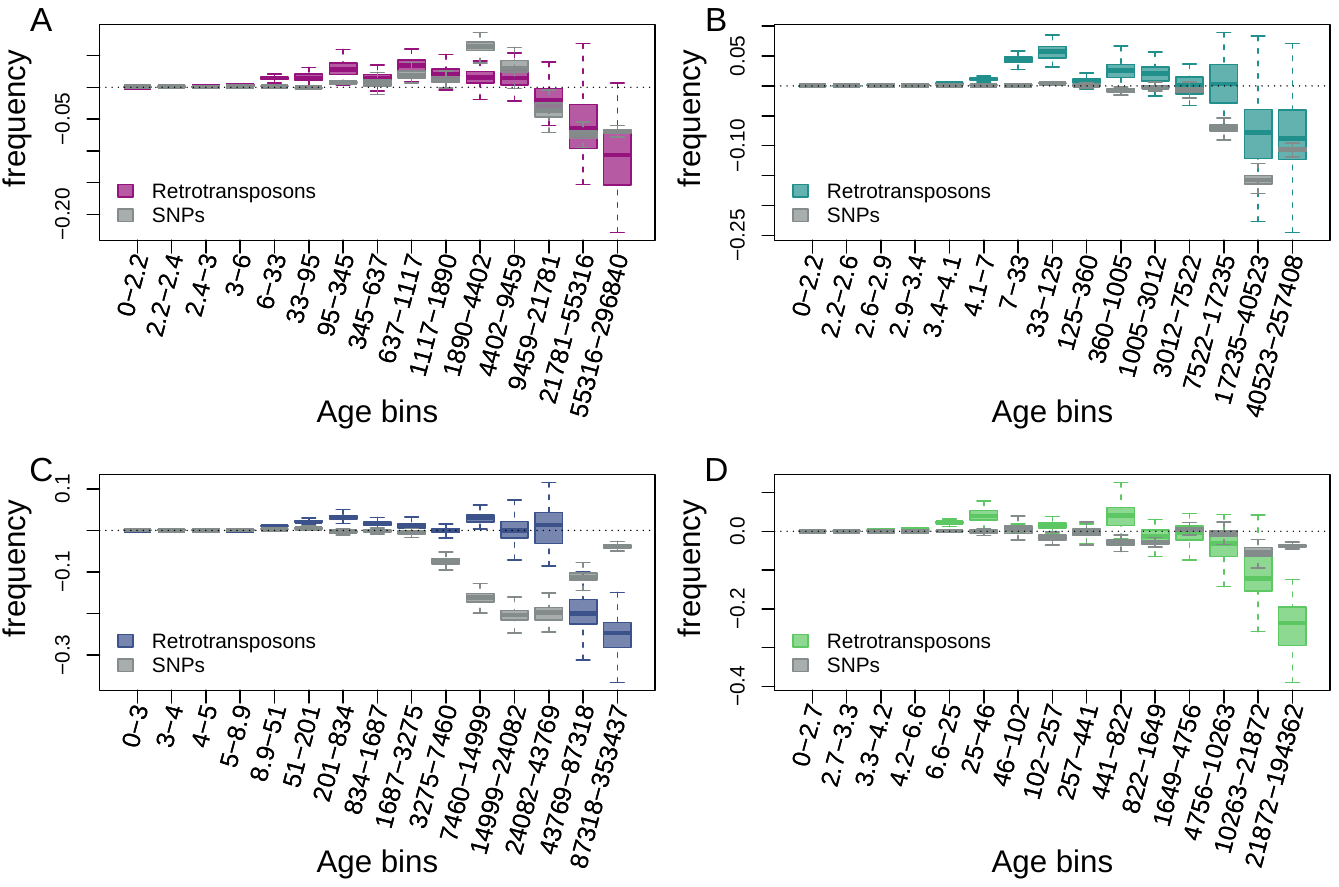

**Supplementary Figure S10.** Age-adjusted SFS of retrotransposons (colored) and SNPs (gray) more than 5 kb away from genes in the four derived clades. The X axes show the age range of the mutations in each bin. A: A_East genetic clades; B: A_Italia genetic clades; C: B_West genetic clades; D: B_East genetic clades. Boxplots are based on 100 estimations of Δ frequency.

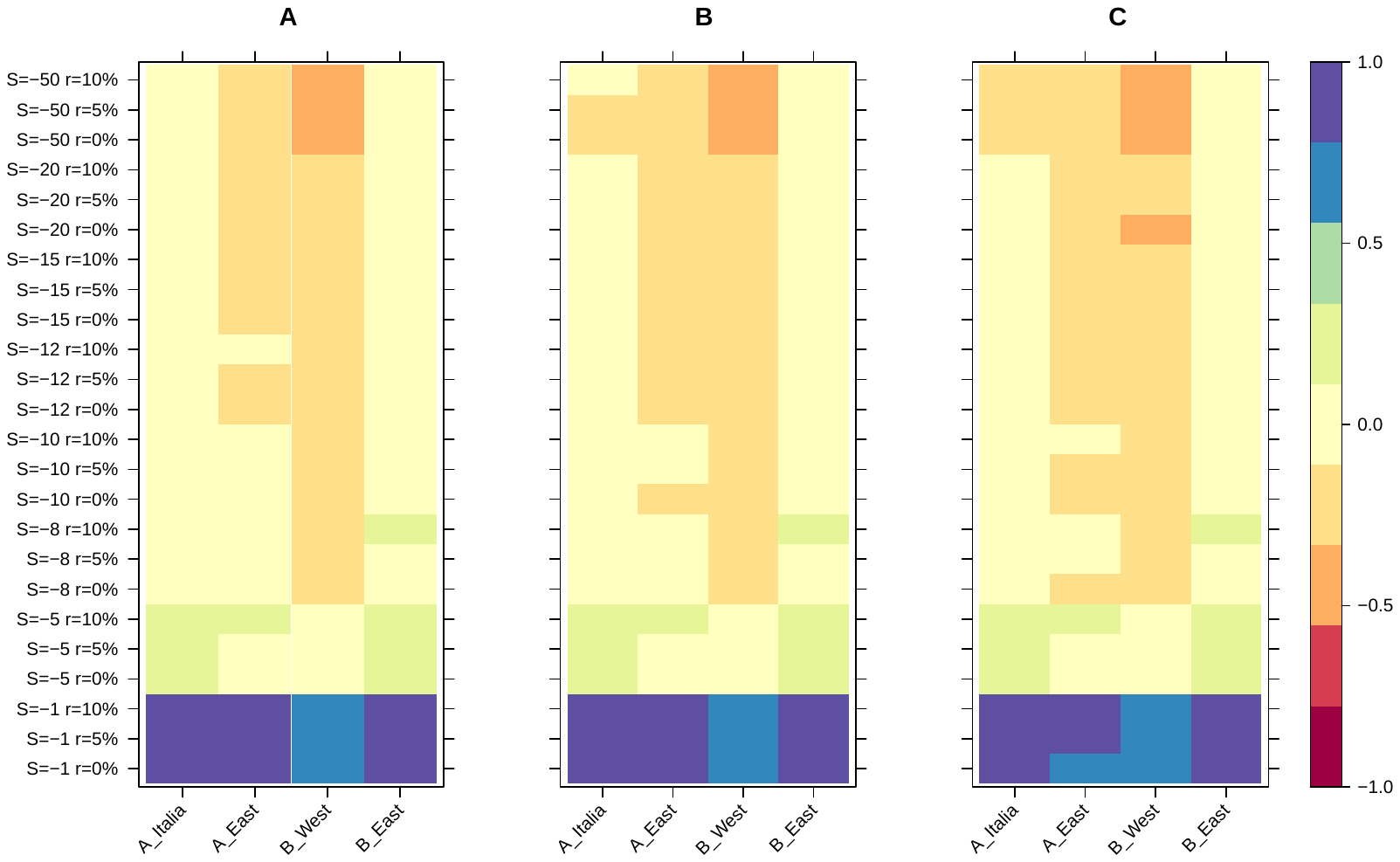

**Supplementary Figure S11.** Relative age difference ((mutation age in simulations - observed mutation age)/maximum absolute age difference) between simulated data assuming fully outcrossing individuals and observed data in the last bin of the age-adjusted SFS. A: 25% quantile; B: 50% quantile; C: 75% quantile.

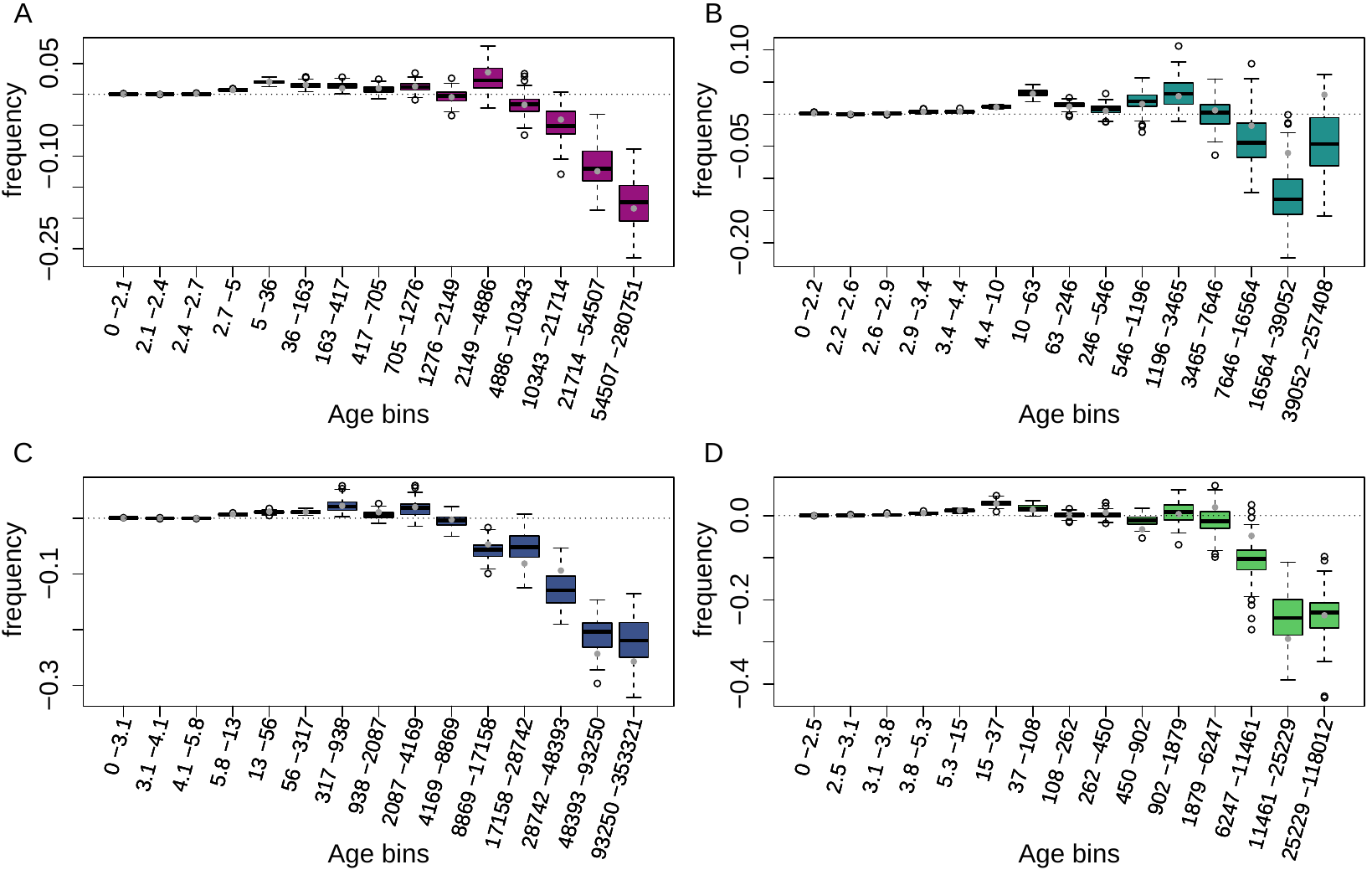

**Supplementary Figure S12.** Age-adjusted SFS of Copia TEs in the four derived clades. The X axes show the age range of the mutations in each bin. A: A_East genetic clades; B: A_Italia genetic clades; C: B_West genetic clades; D: B_East genetic clades. Boxplots are based on 100 estimations of Δ frequency.

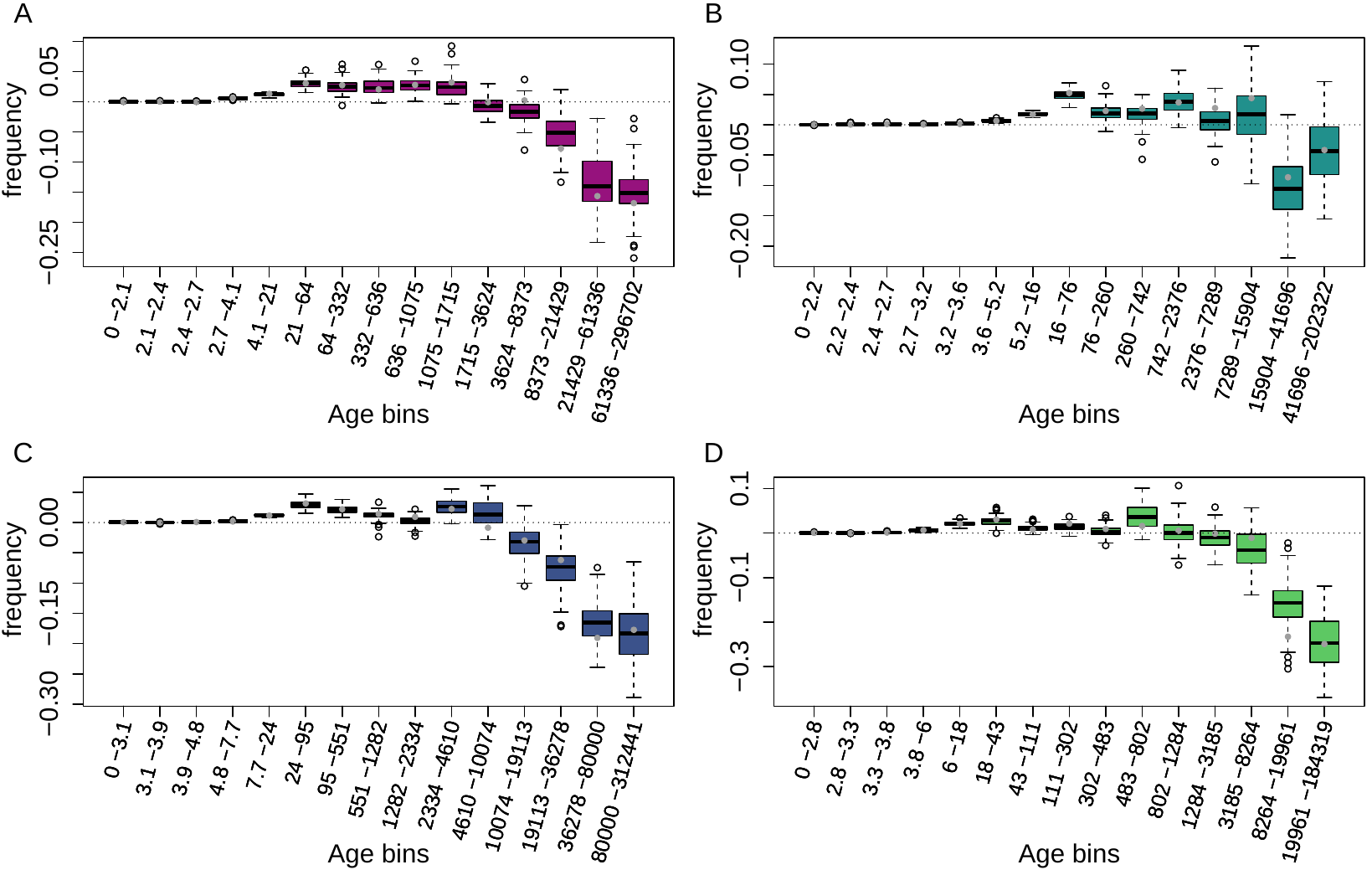

**Supplementary Figure S13.** Age-adjusted SFS of Ty3 TEs in the four derived clades. The X axes show the age range of the mutations in each bin. A: A_East genetic clades; B: A_Italia genetic clades; C: B_West genetic clades; D: B_East genetic clades. Boxplots are based on 100 estimations of Δ frequency.

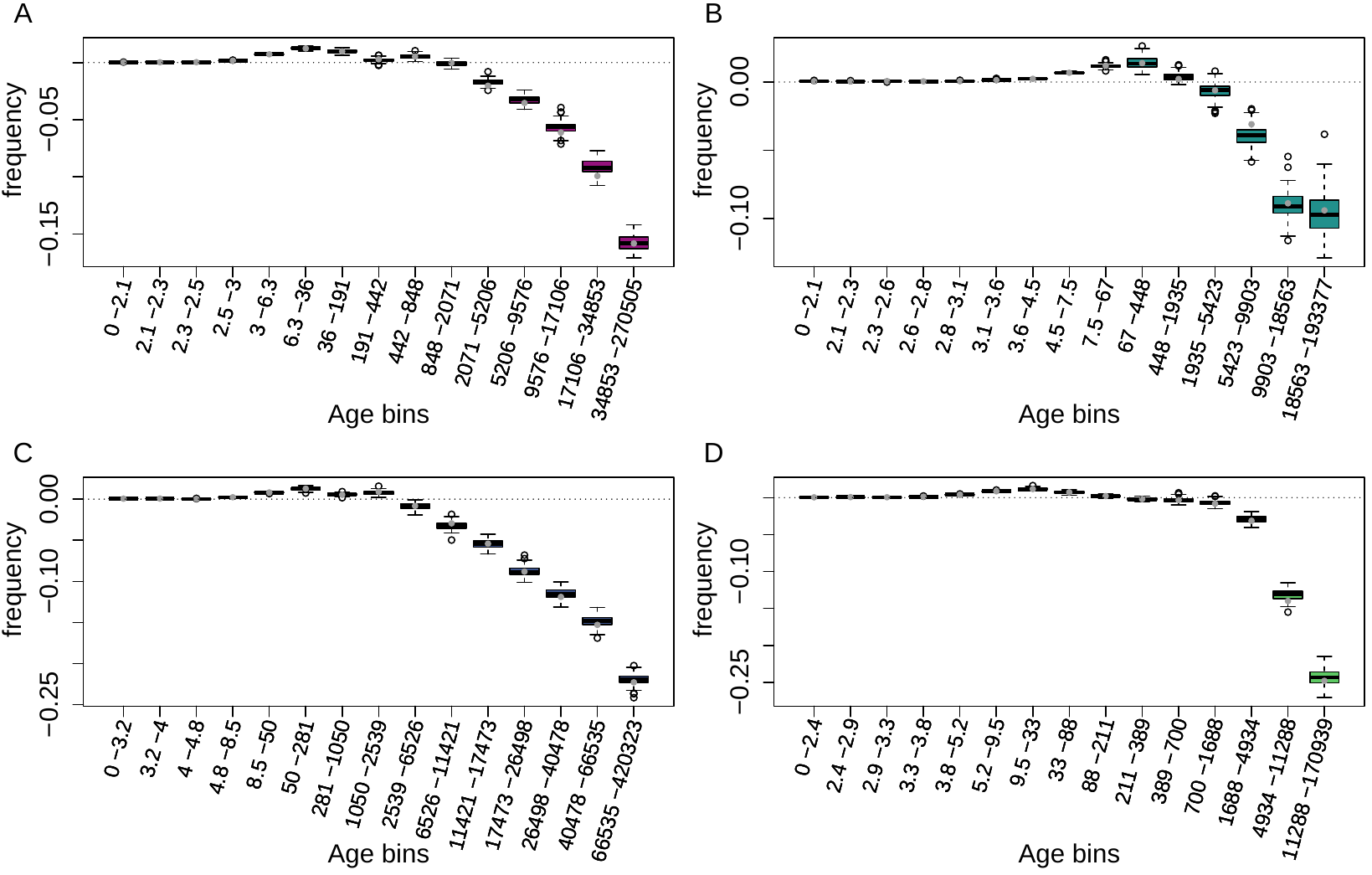

**Supplementary Figure S14.** Age-adjusted SFS of Helitron TEs in the four derived clades. The X axes show the age range of the mutations in each bin. A: A_East genetic clades; B: A_Italia genetic clades; C: B_West genetic clades; D: B_East genetic clades. Boxplots are based on 100 estimations of Δ frequency.

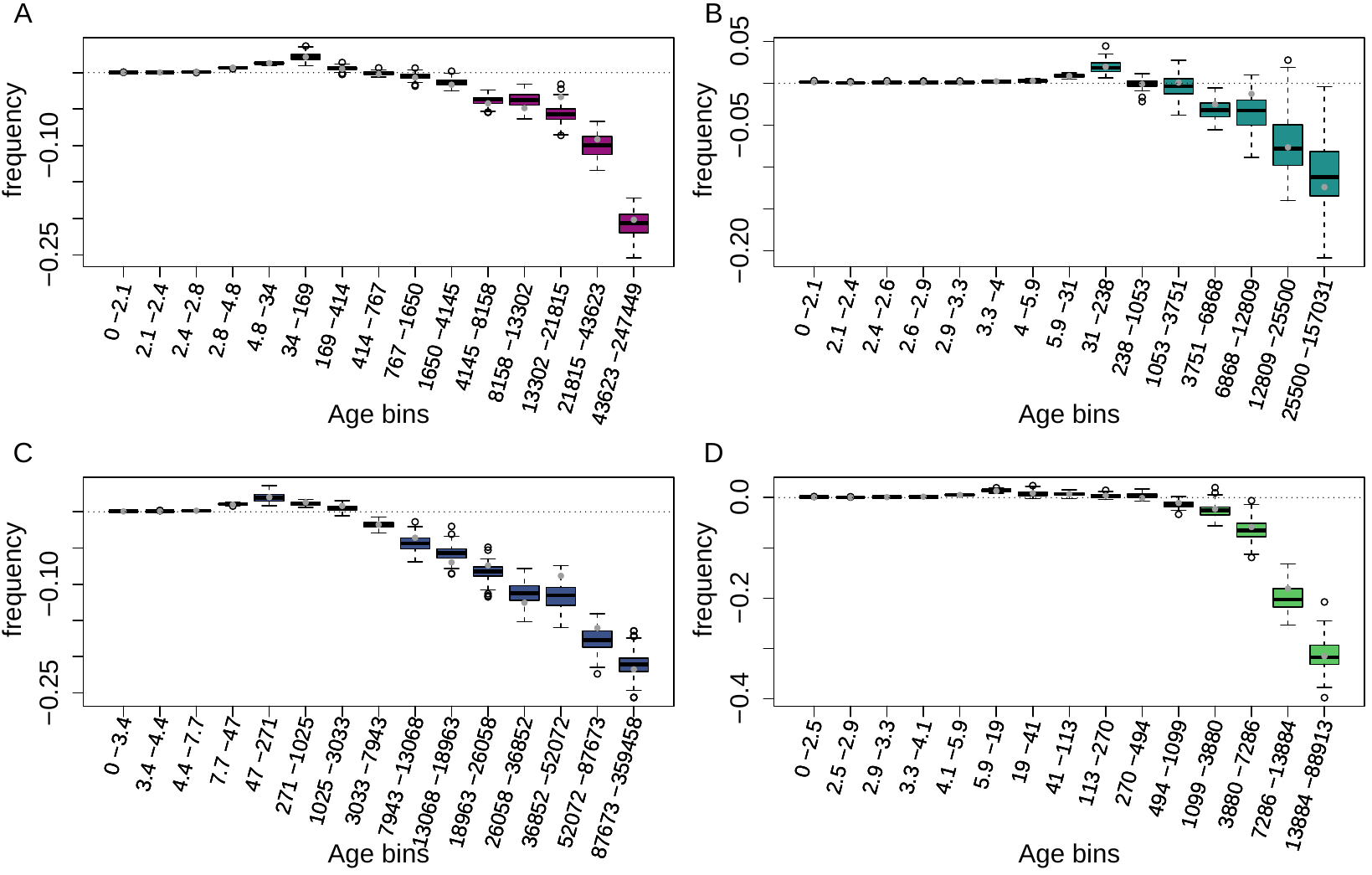

**Supplementary Figure S15.** Age-adjusted SFS of MITE TEs in the four derived clades. The X axes show the age range of the mutations in each bin. A: A_East genetic clades; B: A_Italia genetic clades; C: B_West genetic clades; D: B_East genetic clades. Boxplots are based on 100 estimations of Δ frequency.
